# Delayed exposure of rice to light partially phenocopies C_4_ bundle sheath characteristics by reducing cell length

**DOI:** 10.1101/2023.02.07.527457

**Authors:** Andrew R.G. Plackett, Julian M. Hibberd

## Abstract

C_4_ photosynthesis has arisen in over sixty lineages of angiosperms. Despite significant variation in characteristics of the C_4_ leaf, in all cases a compartment rich in chloroplasts allows high concentrations of CO_2_ to be supplied to RuBisCO. In most C_4_ lineages this is associated with increased chloroplast content of bundle sheath cells. As characteristics of C_4_ leaves are derived from processes operating in the ancestral C_3_ state, we studied greening of the bundle sheath in C_3_ rice. Total visible chloroplast area in bundle sheath and mesophyll cells was similar but the larger volume of bundle sheath cells reduced the proportion of cell occupied by chloroplast, and the longest bundle sheath cells had the lowest chloroplast content. Application of exogenous cytokinin and gibberellin increased chloroplast content of the longest bundle sheath cells but did not change cell shape. In contrast, delayed exposure to light reduced the number of very long bundle sheath cells but increased leaf length indicating delayed exit from the cell cycle. We conclude that in the C_3_ state, the plant hormones cytokinin and gibberellin are regulators of the bundle sheath cell-chloroplast relationship and that final bundle sheath length is affected by light-mediated control of exit from the cell cycle.

**Highlight:** Extended darkness can reduce maximum bundle sheath cell length in rice and hormones condition the set-point between cell size and chloroplast content.

## INTRODUCTION

Photosynthesis in the majority of land plants uses biochemical pathways that operate in algae and photosynthetic bacteria. In these species, the first product of carbon fixation is the three-carbon molecule phosphoglyceric acid and so they have been termed C_3_ plants (Calvin and Benson, 1948; 1949; Benson *et al*., 1950). Synthesis of phosphoglyceric acid is mediated by carboxylase activity of the enzyme Ribulose Bisphosphate Carboxylase Oxygenase (RuBisCO). From around 30 million years ago, species evolved to supplement this ancient biochemistry with an additional pathway involving initial fixation of bicarbonate by the enzyme phospho*enol*pyruvate carboxylase, followed by release of CO_2_ at high concentrations in a compartment containing RuBisCO (Sage *et al*., 2011; Christin *et al*., 2013). In all cases, release of CO_2_ around RuBisCO is mediated by synthesis of C_4_ acids followed by the action of C_4_ acid decarboxylases. This biochemical system serves to repress the competing oxygenase activity of RuBisCO (Leegood, 2002; Carmo-Silva *et al*., 2015) and is known as the C_4_ pathway.

Despite the apparent complexity of C_4_ photosynthesis, which involves the reprogramming of leaf development, alterations to cell ultrastructure and modifications to photosynthetic biochemistry (Langdale, 2011), this trait has evolved independently at least 67 times in the flowering plants (Sage *et al*., 2011; Sage, 2016). In most cases, the reactions of photosynthesis are divided between two distinct cell types with the mesophyll typically becoming specialised for carbon capture whilst the bundle sheath undertakes carbon fixation (Edwards *et al*., 2004). The shuttling of metabolites between these cells is driven by concentration gradients, and so C_4_ leaf anatomy has adapted such that mesophyll cells develop in concentric layers around bundle sheath cells (which in turn develop around veins) and the number of mesophyll cells between veins is reduced to minimise the pathlength for diffusion of the metabolites involved, an adaptation known as ‘Kranz anatomy’ (Sedelnikova *et al*., 2018). In addition, to provide a sufficiently large compartment for CO_2_ fixation by RuBisCO, the size or surface area of C_4_ bundle sheath cells is typically increased compared with C_3_ species (McKown and Dengler, 2007; Christin *et al*., 2013; Lundgren *et al*., 2019, Ermakova *et al*., 2020). C_4_ bundle sheath cells also contain a greatly increased chloroplast content compared with C_3_ plants (McKown and Dengler, 2007; Khoshravesh *et al*., 2016; Wang *et al*., 2017b). Thus, while bundle sheath cells from C_3_ species have a lower chloroplast content than mesophyll cells (Kinsman and Pyke, 1998; Muhaidat *et al*., 2011, Sage *et al*., 2013; Wang *et al*., 2017b; Khoshravesh *et al*., 2020), in C_4_ species the bundle sheath chloroplast content is typically greater than that of the mesophyll. How the bundle sheath of C_4_ plants becomes ‘greener’ compared with that of C_3_ species is poorly understood. One approach to better understand this evolutionary transition would be to investigate the control of chloroplast biogenesis in the C_3_ bundle sheath.

Conceptually, the proportion of cell occupied by chloroplast could be influenced by cell size, itself determined by the balance between cell division and cell expansion (Sablowski, 2016; Jones *et al*., 2019), and within those developing cells the processes of chloroplast biogenesis and chloroplast division (Jarvis and López-Juez, 2013). In the dicotyledenous model *Arabidopsis thaliana* leaf development progresses through co-ordinated organ-wide cell proliferation (combined division and expansion) followed by cell expansion to determine final organ size (Donnelly *et al*., 1999; Beemster *et al*., 2005). Cell and chloroplast development are considered closely co-ordinated with mature mesophyll cells of *A. thaliana* maintaining similar relative chloroplast contents despite variation in individual cell size (Pyke and Leach, 1992; Pyke, 1999). Despite their morphological differences, recent modelling studies have identified homologies between dicotyledonous and monocotyledonous leaf development (Richardson *et al*., 2021) and in both leaf types chloroplast biogenesis and division are controlled by a partially-conserved network of environmental and endogenous signals interfacing with downstream transcription factors such as members of the *GOLDEN2-LIKE* (*GLK*) and *GATA* families (Naito *et al*., 2007; Chiang *et al*., 2012; Zubo *et al*., 2018; Wang *et al*., 2017b; Lee *et al*., 2021). Light and multiple plant hormones regulate these interacting transcription factors to promote chloroplast development (Richter *et al*., 2013; Cackett *et al*., 2022) and in *A. thaliana* a number of the same phytohormones (auxin, cytokinin [CK] and gibberellin [GA]) also regulate cell division and expansion (Hu *et al*., 2003; Schruff *et al*., 2006; Achard *et al*., 2009; Holst *et al*., 2011; Claeys *et al*., 2012; Jiang *et al*., 2012). There is evidence in the monocotyledonous C_3_ model rice (*Oryza sativa*) that both cell and chloroplast development are also regulated by these signals (Matsukura *et al*., 1998; Ikeda *et al*. 2001, Yang *et al*., 2002, Jiang *et al*., 2012, Aya *et al*., 2014, van Campen *et al*., 2016, Ding *et al*., 2017). However, to our knowledge their roles in bundle sheath cell development have not been investigated.

To test the extent to which these regulatory pathways influence chloroplast content of bundle sheath cells in rice we manipulated hormone and light inputs during leaf development and determined their effect on mature bundle sheath and mesophyll cell anatomy. This indicated that although bundle sheath chloroplast biogenesis increases with cell size a ‘set-point’ relationship between chloroplast content and cell area is not maintained. Exogenous hormone treatments significantly affected the relationship between cell size and chloroplast biogenesis such that larger cells contained more chloroplast material at maturity. Delaying exposure of leaves to light changed the shape of mature bundle sheath cells by reducing length. As total leaf length was simultaneously increased this indicates that light promotes exit of bundle sheath cells from the cell cycle. We conclude that hormone signalling co-ordinates chloroplast biogenesis and bundle sheath cell size, and propose that enhanced greening of bundle sheath cells in C_4_ grasses has likely arisen through re-wiring of these ancestral light- and hormone-dependent signalling pathways.

## MATERIALS AND METHODS

### Plant material and growth conditions

All experiments were performed using rice (*Oryza sativa*) cv. Kitaake. Seed was imbibed in sterile water and germinated on filter paper in darkness at 32°C for three days before transplanting to soil. Germinating seeds were sown into a 1:1 mix of Erin topsoil (LBS Horticulture, Colne, UK) and washed sand, supplemented with 1x Everris Peters Excel Cal-Mag Grower fertilizer solution (LBS Horticulture, Colne, UK) with additional 0.35% chelated iron (w/v) (Garden Direct, Stevenage, UK). Seeds were sown into 7 cm pots and kept covered with transparent propagator lids for four days after sowing. All plants were grown in a controlled environment growth room under a photoperiod of 12 hours light and 12 hours dark, a temperature of 28°C (day) and 25°C (night), a constant relative humidity of 60% and a photon flux density of 300 μmol m^-2^ s^-1^.

### Exogenous hormone treatment

Exogenous hormone treatments comprised 5 μM 1-napthaleneacetic acid (NAA; Merck Life Science UK Ltd., Gillingham, UK) (0.1% v/v Tween20), 5 μM 6-benzylaminopurine (BAP; Merck Life Science UK Ltd., Gillingham, UK) (0.1% v/v Tween20), 100 μM gibberellin A3 (GA_3_; Melford Laboratories Ltd., Ipswich, UK) (0.1% v/v ethanol, 0.1% v/v Tween20) and mock solution (0.1% v/v ethanol; 0.1% v/v Tween20). Hormone treatments were applied to whole plants as a foliar spray, taking care to coat the entire plant and supply an equal quantity of hormone solution (5 sprays per plant). Hormone solutions were stored at 4°C and brought to room temperature before use. Plants were treated every 2-3 days from four days after sowing onwards. Measurements of leaf 4 length were taken at 15 days after sowing, measuring six plants per treatment. On the same day leaf 4 tissue was harvested from the same plants for single cell isolation - 5 mm of blade tissue either side of the blade mid-point was taken. Leaf tissue was imaged from three plants per treatment, and from five mesophyll and five bundle sheath cells per plant.

### Delayed light exposure

Imbibed seed was sown to soil in a darkroom under green light at a density of three seeds per pot. Light-grown control plants were incubated under a transparent propagator lid for four days, whereas plants under dark-grown and delayed-light treatments were grown in the same conditions whilst avoiding any exposure to light. Delayed-light plants were exposed to standard light-grown conditions six days after sowing. Dark-grown plants were exposed to standard light-grown conditions immediately before tissue harvesting (eight days after sowing). Leaf 2 blade length was measured from six plants per treatment. Leaf blade tissue was harvested for single cell isolation on the same day, taking 5 mm either side of the blade mid-point. Leaf tissue was imaged from six plants per treatment, and from five mesophyll and five bundle sheath cells per plant. To determine dark-grown leaf 2 growth responses to light exposure, 48 plants were grown under dark-grown treatment as above. At seven days after sowing 24 plants were exposed to standard light-grown conditions and leaf 2 blade length was measured at 0, 0.5, 1, 2, 3, 4, 6 and 24 hours after light exposure. Leaf 2 blade length was measured from three light-induced and three corresponding plants not exposed to light at each timepoint. For consistency, three independent plants were measured per timepoint under each treatment.

### Premature exposure to light

Imbibed seeds were sown at a density of three seeds per pot and exposed to standard light-grown conditions from seed sowing onwards. Premature exposure of leaf 4 to light was achieved through the manual removal of the surrounding leaf 3. On the day that the tip of the leaf 4 primordium emerged from within the sheath of leaf 3 (typically 6-8 days after sowing), leaf 3 was removed by initially tearing the blade in half lengthways (along the proximodistal axis) using two pairs of watchmakers’ forceps and then extending the tear down through the sheath to as close to the plant base as possible. Removal of only the leaf 3 blade was achieved by cutting with scissors at or just above the leaf 3 ligule. For leaf 4 length measurements, one replicate of each treatment (control, leaf 4 exposure, leaf 3 blade removed) was applied per pot on the day of leaf 4 emergence, and leaf 4 length was subsequently measured every 24 hours until leaf growth had ceased in all plants. Leaf 4 length was measured from 20 replicates per treatment. For leaf 4 cell measurements, leaf 4 tissue was harvested three days after premature exposure and simultaneously from control plants within the same population, taking 5 mm of blade tissue above the ligule. Leaf tissue was imaged from three plants per treatment, and from ten mesophyll, ten bundle sheath and ten epidermal cells per plant.

### Single cell isolation and imaging

Rice leaf cells were isolated following the protocol of Khoshravesh and Sage (2018). Leaf tissues were cut into 5 mm-long x 2 mm-wide strips along the leaf proximodistal axis in a drop of water using a razorblade and immediately immersed in room temperature 4% w/v paraformaldehyde (pH6.9) (Merck Life Science UK Ltd., Gillingham, UK). Tissue was immediately placed in darkness and fixed at 4°C overnight, after which fixed tissue was stored in 1x PBS solution (pH7.0) at 4°C. Cell walls were digested by incubating in 0.2M sodium-EDTA (pH9.0) 55°C for 2 hours, then 2% w/v *Aspegillus niger* pectinase (Merck Life Science UK Ltd., Gillingham, UK) 45°C for 2 hours. Digestion was stopped by incubation in empty buffer twice for 30 minutes at room temperature. Individual cells were imaged within 24 hours of cell wall digestion.

Isolated cells were imaged by brightfield microscopy using an Olympus BX51 microscope (Olympus UK and Ireland, Southend-on-Sea, UK), recording each cell in the paradermal plane at 4-5 separate focal depths, focussed on the varying distribution of chloroplasts within the cell. Images were captured using an MP3.3-RTV-R-CLR-10-C MicroPublisher camera and QCapture Pro 7 software (Teledyne Photometrics, Birmingham, UK). Only bundle sheath cells still attached to vascular tissue were selected for imaging. Cell and chloroplast measurements were taken from scaled images using FIJI (Schindelin *et al*., 2012), using the multiple images captured per cell to increase the accuracy of measurement at the whole-cell level. Whole plant photographs were taken using a Cybershot DSC-HX7V digital camera (Sony, Tokyo, Japan). Figures were prepared using Photoshop 23.0.2 (Adobe, San Jose, USA). Photographic images included in these were individually adjusted for brightness and contrast.

### Data analysis

Statistical analyses were performed in RStudio version 1.2.5033 (RStudio Team, 2020) using R version 3.6.3 (R Core Team, 2020). ANOVA and regression analyses were undertaken using the car software package (Fox and Weisberg, 2019). Relative chloroplast content was calculated by dividing total chloroplast area by the corresponding paradermal cell area. The range of cell lengths within a cell type + treatment combination was normalised relative to the smallest cell within that category. As an estimate of chloroplast size the largest chloroplast visible per cell was chosen as the least-biased method of comparison. Datasets were tested for Normal distributions using the Shapiro-Wilk test (Shapiro and Wilk, 1965) and variances were compared using Levene’s test (Levene, 1960). Where the assumptions of ANOVA were met pairwise comparisons were made using two-tailed pairwise T-tests, otherwise they were made using two-tailed Mann-Whitney tests. Where multiple pairwise comparisons were made within an experiment, the p-values obtained were corrected post-hoc to minimise false discoveries (Benjamini and Hochberg, 1995). All cell measurements, transformations and statistical test outputs are given in **Supplementary Datasets S1-S4**. All plots were prepared with the ggplot2 software package (Wickham, 2016).

## RESULTS

### Chloroplast biogenesis responds to size of mesophyll and bundle sheath cells but a set point is not maintained in the bundle sheath

To better understand developmental processes associated with differences in chloroplast content between rice mesophyll and bundle sheath cells we first characterised these cell types at maturity (**Fig. 1A, B**). As expected, the area of cell occupied by chloroplasts (hereafter relative chloroplast content) was significantly lower for bundle sheath than mesophyll cells (p < 0.05; **Fig. 1C**). Although no statistically significant difference was detected between the total area of chloroplast per mesophyll or bundle sheath cell (p > 0.05, **Fig. 1D**) both the number and size of chloroplasts was lower in bundle sheath cells (p < 0.05, **Fig. 1E, F**). We note that the two-dimensional method of image analysis employed here could underestimate chloroplast area in mesophyll cells, where chloroplasts can overlap (**Fig. 1A**). Bundle sheath cell area and cell length parallel to the leaf proximodistal axis (along the vein) were greater than mesophyll cells (p < 0.05), whereas cell width (parallel to the leaf mediolateral axis) was smaller (p < 0.05; **Supplementary Dataset S1**). Analysis of variance indicated that measurements from individual cells were statistically independent of the plant from which they were isolated (p > 0.05; **Supplementary Dataset S1**) and so each cell could be treated as an independent replicate. Bundle sheath cells demonstrated greater variance in cell area and length than mesophyll cells (p < 0.05, Levene’s test), but variance in cell width was similar (p = 0.055). Similarly, except for size, chloroplast characters of bundle sheath cells (including relative chloroplast content) had greater variance than those of mesophyll cells (p < 0.05). These results confirmed that individual cells within the bundle sheath are more variable in their morphology than mesophyll cells, and that relative chloroplast content of the bundle sheath is reduced compared with mesophyll cells. Our results suggest that both increased cell size and reduced chloroplast biogenesis (smaller, fewer chloroplasts) in bundle sheath cells contribute to the lower relative chloroplast content compared with the mesophyll.

**Fig. 1.**
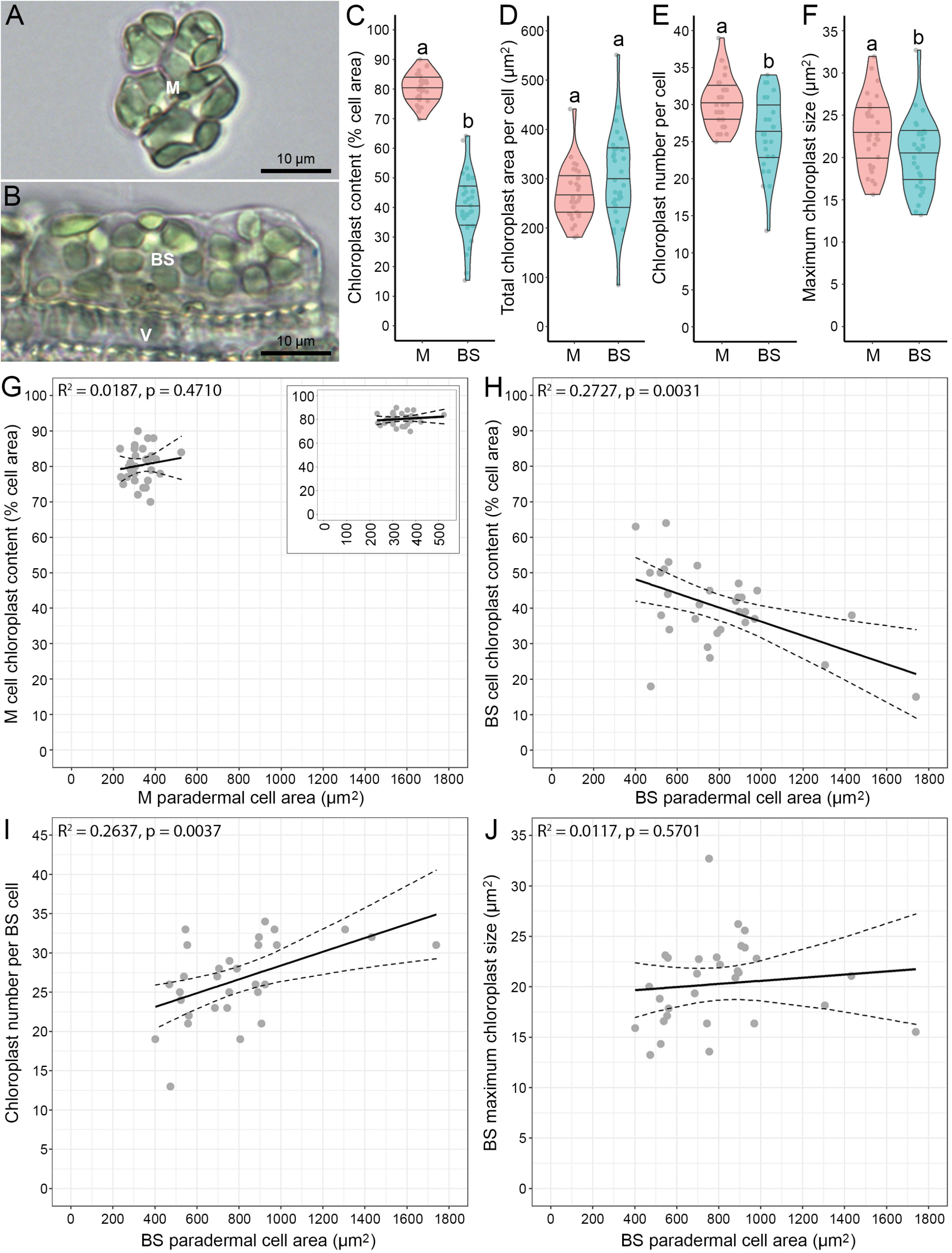
Chloroplast biogenesis is reduced in rice bundle sheath compared with mesophyll cells. **A-B**. Representative images of individual mesophyll (**A**) and bundle sheath (**B**) cells isolated from mature leaf tissue. Images are oriented to follow the leaf proximodistal axis (left to right). M, mesophyll cell; BS, bundle sheath cell; V, vascular bundle. Bundle sheath cells remained attached to the vascular bundle after partial digestion. **C-F**. Comparison of relative chloroplast content (**C**), total chloroplast area (**D**), chloroplast number (**E**) and maximum chloroplast size (**F**) per cell between mature mesophyll (M) and bundle sheath (BS) cells isolated from leaf 4. Violin plots represent data distribution density, with their width corresponding to frequency of datapoints (grey circles) at that value. Median value and upper and lower quartiles are represented by the middle, upper and lower horizontal lines, respectively. Different letters denote statistically significant differences (p < 0.05) between BS and M cells within each plot. Comparisons of relative chloroplast content and chloroplast size were made using transformed datasets to meet the assumptions of statistical tests (see **Supplementary Dataset S1**). **G-J**. Linear regression analyses of relative chloroplast content against cell area for M (**G**) and BS cells (**H**), and of BS cell chloroplast number (**I**) and size (**J**) against cell area, respectively. Graphs show the line of best fit and 95% confidence interval of the linear model fitted to the data (grey circles). Insert in (**G**) shows same data with x-axis re-scaled to better visualise the distribution of cell areas captured. *n* = 30 (5 cells of each type measured from 6 individual plants, see **Supplementary Fig. S1A-F**).

To investigate the relationship between cell and chloroplast development in more detail, regression analysis was used (**Fig. 1G-J**). For mesophyll cells, no significant relationship was detected between relative chloroplast content and cell area (p = 0.4710; **Fig. 1G**). In contrast, for the bundle sheath a significant negative relationship was apparent (p = 0.0031; **Fig 1H**). This is unlikely to be an artefact caused by swelling of cells during digestion of the cell wall because the same relationship was detected between relative chloroplast content and cell length (p = 0.0010; **Supplementary Dataset S1**). Length of bundle sheath cells has limited capacity to alter as it is constrained by attachment to the vascular bundles (**Fig. 1A**). No statistically significant relationship was detected between bundle sheath chloroplast size and cell area (p = 0.5701; **Fig. 1J**) but robust positive relationships were found between cell area and chloroplast number (p = 0.0037, **Fig. 1I**) and between cell area and total chloroplast area per cell (p = 0.0003; **Supplementary Dataset S1**). These two positive relationships were also detected for mesophyll cells (p < 0.05), as was a marginal positive relationship between chloroplast size and cell area (**Supplementary Fig. S1A, B; Supplementary Dataset S1**). No statistically significant relationship was found between chloroplast size and chloroplast number for either cell type (**Supplementary Dataset S1**). Measurements of relative chloroplast content from individual bundle sheath and mesophyll cells were not statistically independent of the plant from which they were isolated (p = 0.0129 and p = 0.0483 respectively) and inclusion of this parameter increased explanatory power of linear models (**Supplemental Fig. S1C, D**) but did not significantly affect the relationship between chloroplast content and cell area (**Supplementary Dataset S1**). The plant from which cells were isolated was also a significant explanatory variate for chloroplast size but not chloroplast number in both the bundle sheath and mesophyll (**Supplementary Fig. S1E, F**; **Supplementary Dataset S1**). Mindful of the limitations of two-dimensional cell analysis, we tested the effect of this on our interpretation of changing chloroplast number with cell size. As published transverse sections that demonstrate rice bundle sheath cells are approximately circular in cross-section (Wang *et al*., 2017b) we modelled individual bundle sheath cells as cylinders using cell length and width measurements (**Supplementary Fig. S2A-D**). Chloroplast number still showed a positive relationship with cell volume (p = 0.0012; **Supplementary Fig. S2E**) supporting the robustness of this relationship. Overall, these results indicate that chloroplast biogenesis responds to increasing cell size in the bundle sheath, but in contrast to the mesophyll this response is not sufficiently strong to maintain a constant chloroplast content. We conclude that mechanisms coordinating cell and chloroplast development are active in both the mesophyll and bundle sheath cells, but they are differentially regulated between the two cell types.

### Cytokinin and gibberellin increase chloroplast biogenesis in the bundle sheath in a cell size dependent manner

The ability of cytokinin, auxin and GA to affect cell and chloroplast development in rice was tested by exogenous application of each hormone during development of leaf 4. Final length of leaf 4 was not altered by treatment with cytokinin (BAP) or auxin (NAA) (p > 0.05) but in gibberellin (GA)-treated plants leaf blade and leaf sheath length was increased compared with mock-treated controls (p < 0.05; **Fig. 2A**). The proportion of blade to sheath was significantly reduced by GA treatment (p < 0.05) (**Supplementary Dataset S2**). None of the hormones had a detectable impact on paradermal area of mesophyll or bundle sheath cells (p > 0.05; **Fig. 2B**), or bundle sheath cell length along the leaf proximodistal axis (**Supplementary Dataset S2**). As GA application generated a longer leaf blade but cell length was not changed, cell division in the leaf blade must have been prolonged by this treatment.

**Fig. 2.**
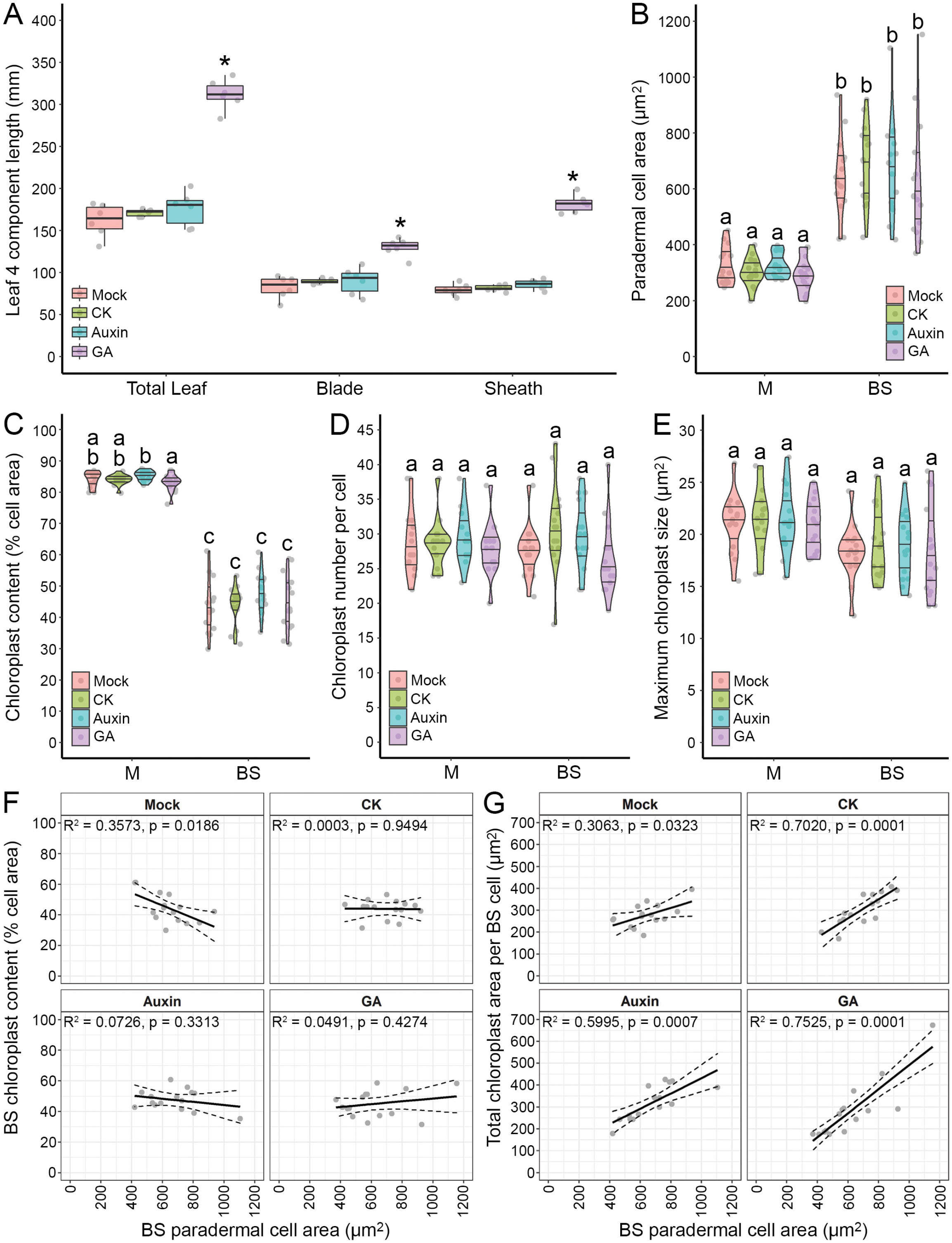
Exogenous hormone treatments alter rice BS cell-chloroplast relationships. **A**. Comparison of mature leaf 4 length characteristics between mock (0.1% EtOH v/v), 5 μM BAP, 5 μM NAA and 100 μM GA_3_ treatments (as shown). Boxplots represent the 25^th^-75^th^ percentile (box), median value (mid-line) further datapoints within 1.5x of the interquartile distance (whiskers). Asterisks denote significant difference (p < 0.05) from mock treatment. *n* = 6. **B-E**. Comparison of bundle sheath (BS) and mesophyll (M) cell paradermal area (**B**), relative chloroplast content (**C**), chloroplast number per cell (**D**) and maximum chloroplast size (**E**) from mature leaf 4 between mock (0.1% EtOH v/v), 5 μM BAP, 5 μM NAA and 100 μM GA_3_ treatments (as shown). Violin plots represent data distribution density, with their width corresponding to frequency of datapoints (grey circles) at that value. Median value and upper and lower quartiles are represented by the middle, upper and lower horizontal lines, respectively. Different letters denote a significant difference (p < 0.05) between cell type + hormone treatment combinations within each plot. Comparisons within all characters except chloroplast size were made using transformed datasets to meet the assumptions of statistical tests (**Supplementary Dataset S2**). **F**-**G**. Regression analyses of relative chloroplast content (**F**) and total chloroplast area per cell (**G**) against cell area in BS cells under mock (0.1% EtOH v/v), 5 μM BAP, 5 μM NAA and 100 μM GA_3_ treatments (as shown), showing the line of best fit and 95% confidence interval of the model fitted to the data (grey circles). P-values and R^2^ values are from regression analyses of the data against cell area within each treatment, denoting the significance of cell area as an explanatory variate and the explanatory power of the fitted model, respectively. *n* = 15 (5 cells measured from 3 individual plants).

Hormone treatments did not alter relative chloroplast content (**Fig. 2C**), chloroplast number per cell (**Fig. 2D**) or maximum chloroplast size (**Fig. 2E**) compared with the mock treatment in either cell type. Moreover, for the mesophyll, no significant effect of hormone treatments on the relationship between relative chloroplast content and cell area was detected by regression analysis (p > 0.05, **Supplementary Fig. S3A, B; Supplementary Dataset S2**). However, for the bundle sheath, whilst a negative relationship between cell area and chloroplast content was apparent for each of the mock-treatments (p < 0.05 and consistent with data in **Fig. 1H**), when auxin, CK or GA were applied this relationship was no longer statistically significant (**Fig. 2F**). Pairwise comparison of each hormone treatment against the mock confirmed that the relationship between relative chloroplast content and cell area was altered by each hormone to become more positive (p < 0.05 and **Supplemental Fig. S3C-E**; **Supplementary Dataset S2**). Moreover, while mock and hormone-treated plants all demonstrated a weak significant positive relationship between total chloroplast area and bundle sheath cell area (p < 0.05, **Fig. 2G**) this correlation was strengthened by each hormone: notably, CK and GA had statistically significant effects on the relationship between cell area and total chloroplast content (p < 0.05) (**Supplementary Fig. S3F-H**). The effects remained robust when potential outlier datapoints were excluded (**Supplementary Dataset S2**). These data indicate that treatment with CK and GA enhanced chloroplast biogenesis in larger bundle sheath cells, and so we conclude they impact on the relationship between chloroplast content and cell development of the rice bundle sheath.

### Darkness reduces bundle sheath length

As hormones and light signalling interact to control chloroplast biogenesis we next tested the effect of light signalling on cell and chloroplast development in the rice bundle sheath. In control experiments leaf 2 completed development by seven days after germination (**Fig. 3A**) at which time growth of its blade had ceased whether exposed to light or not (**Supplementary Fig. S4A**). In addition to comparing cell morphology from light or dark grown plants, a treatment in which exposure to light was delayed was provided. Delayed light exposure altered final leaf length but also bundle sheath cell length. Mature blade length was significantly increased in both dark-grown and delayed-light plants compared with light-grown controls (p < 0.05; **Fig 3B**). Green chloroplasts were visible in bundle sheath and mesophyll cells of light-but not dark-grown plants (**Fig. 3C, 3D, 3F, 3G**), whereas in delayed-light plants chloroplasts from both cell types had only partially greened (**Fig. 3E, 3H**). In contrast with the increase in leaf length, the mean area of mesophyll and bundle sheath cells was significantly reduced in dark-grown leaves compared with light-grown controls (p < 0.05 **Fig. 3I**). Interestingly, under delayed light while mesophyll cell area was similar to dark-grown cells (p > 0.05), bundle sheath cell area was intermediate between dark-grown and light-grown controls (**Fig. 3I**). Thus, leaf 2 responded to the delayed light signal and this modified bundle sheath morphology.

**Fig. 3.**
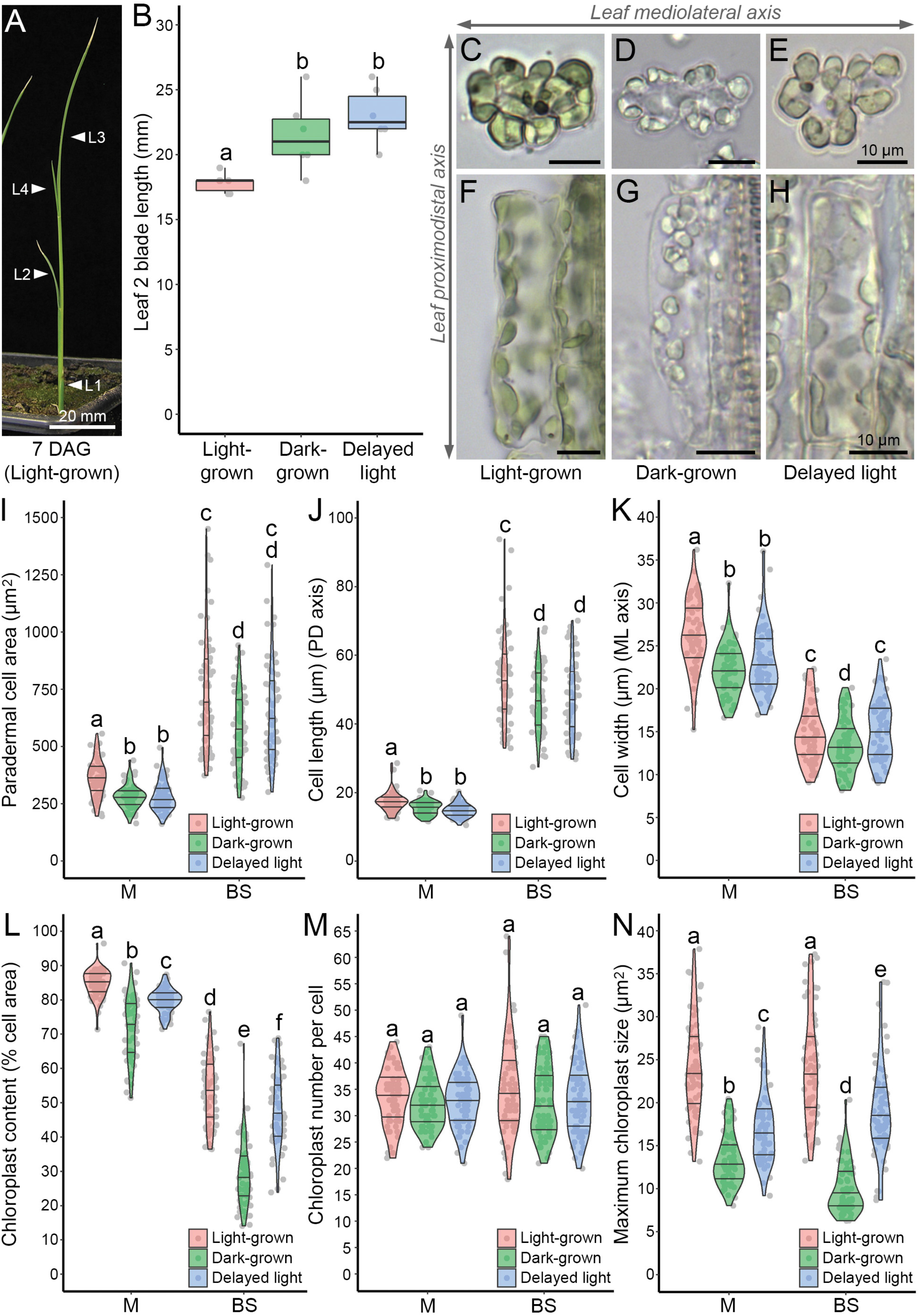
Delayed light exposure prolongs cell division of photosynthetic cells in the rice leaf. **A**. WT rice plant (cultivar Kitaake) 7 days after germination (DAG) under light-grown conditions. The relative position of the four visible leaves (L1-L4) are indicated (arrowheads). **B**. Comparison of leaf 2 blade length between 8 day-old light-grown, dark-grown and delayed light treatments (as shown). Boxplots represent the 25^th^-75^th^ percentile (box), median value (mid-line) further datapoints within 1.5x of the interquartile distance (whiskers). Different letters denote statistically significant differences (p < 0.05) between light treatments. *n* = 6. **C-H**. Experimental light treatments caused visible changes in chloroplast morphology of both BS (**C-E**) and M cells (**F-H**). Compared to cells from 8 day-old control plants exposed to light from germination onwards (‘Light-grown’; **C, F**), cells from 8 day-old plants not exposed to light (‘Dark-grown’) contained etioplasts visibly lacking chlorophyll (**D, G**). Cells from 8 day-old plants exposed to delayed light at 6d after germination (‘Delayed light’) exhibited visible greening compared to dark-grown controls but not to the extent of light-grown controls (**E, H**). Cell images are shown oriented with respect to the axes along the length of the leaf blade (proximodistal) and across its width (mediolateral). **I-N**. Comparison of cell paradermal area (**I**), length (**J**), width (**K**), relative chloroplast content (**L**), chloroplast number per cell (**M**) and maximum chloroplast size (**N**) of mesophyll (M) and bundle sheath (BS) cells isolated from the 8 day-old leaf 2 blade between light-grown, dark-grown and delayed light treatments (as shown). Violin plots represent data distribution density, with their width corresponding to frequency of datapoints at that value. Median value and upper and lower quartiles are represented by the middle, upper and lower horizontal lines, respectively. Different letters denote a significant difference (p < 0.05) between cell type + light treatment combinations within each plot. Comparisons within all characters except leaf length were made using transformed datasets to meet the assumptions of statistical tests (see **Supplementary Dataset S3**). *n* = 60 (10 cells of each type measured from 6 individual plants per treatment).

The perturbation to bundle sheath phenotype under delayed light indicates that both cell expansion and cell division are light regulated in this tissue. Length and width of mesophyll cells were reduced by both the dark-grown and delayed light treatments compared with light-grown controls (p < 0.05) (**Fig. 3J, K**). However, for the bundle sheath while cell length was reduced by both dark treatment and delayed exposure to light (p < 0.05; **Fig. 3J**), cell width was only reduced in dark-grown plants (p < 0.05; **Fig. 3K**). The reduced length of bundle sheath cells along the leaf proximodistal axis in dark-grown and delayed light treatments was associated with absence of the longest cells (**Fig. 3J)**. For example, in light-grown controls maximum bundle sheath cell length was 2.9 times that of the shortest cell, but under dark-grown and delayed-light treatments this parameter was reduced to 2.5 and 2.4 respectively (**Supplementary Dataset S3**). Variance in bundle sheath cell lengths was similar between all three treatments (p > 0.05, Levene’s test), but while in light-grown controls the longest cells meant that the data were not Normally distributed (p = 0.007, Shapiro-Wilk test), in dark-grown and delayed-light populations they were (p = 0.228 and p = 0.067 respectively). Chloroplast content of both mesophyll and bundle sheath cells from leaves subjected to delayed light was intermediate between light-grown and dark-grown controls and significantly different from both (p < 0.05; **Fig. 3L**). Chloroplast number per cell was not affected by delayed light or absence of light (p > 0.05; **Fig. 3M**). However, with delayed light chloroplasts were intermediate in size between those from both dark-grown and light-grown controls (p < 0.05, **Fig. 3N**). Our findings that absence of light increased total leaf length (**Fig. 3B**) but reduced mesophyll and bundle sheath cell length (**Fig. 3J**) are most easily rationalised by additional rounds of cell division taking place in the bundle sheath and mesophyll, a hypothesis supported by the changing distribution of cell lengths. The subsequent recovery of bundle sheath cell width when exposed to delayed light suggests that light triggers bundle sheath cell expansion at the same time as photosynthetic activation.

### Premature light reduces leaf but not bundle sheath length

The above results are consistent with light promoting exit of photosynthetic cells in rice from the cell cycle, a function previously proposed in *A. thaliana* (Andriankaja *et al*., 2012). We therefore hypothesised that premature exposure to light would cause earlier exit from the cell cycle and so result in shorter leaves. To test this the primordium of leaf 4 was exposed to light prematurely by removal or leaf 3 (**Fig. 4A-C**) and the length of leaf 4 recorded until full expansion was achieved (**Fig. 4D-F**). For technical reasons leaf 3 could only be removed when the tip of leaf 4 emerged above the leaf 3 ligule (6-8 days after germination, **Fig. 4A, C**). To distinguish between direct effects of light signalling on leaf 4 growth, and indirect effects caused by a reduction in photosynthate supply once leaf 3 was removed, an additional treatment was included whereby the blade of leaf 3 was removed whilst leaf 4 remained in the leaf 3 sheath (**Fig. 4B**). When exposed prematurely to light, leaf 4 length was significantly reduced at maturity compared with controls (p < 0.05) (**Fig. 4G**). Removal of the leaf 3 blade did not affect the final length of leaf 4 (p > 0.05). Premature exposure to light caused a reduction in the rate of leaf 4 growth rather than premature termination of its development (**Supplementary Fig. S4B**). These results strongly support a role for light signalling in regulating leaf length, and also demonstrate that the growth habit in which rice leaf primordia arise within older leaves influences their development through altering the timing of light exposure.

**Fig. 4.**
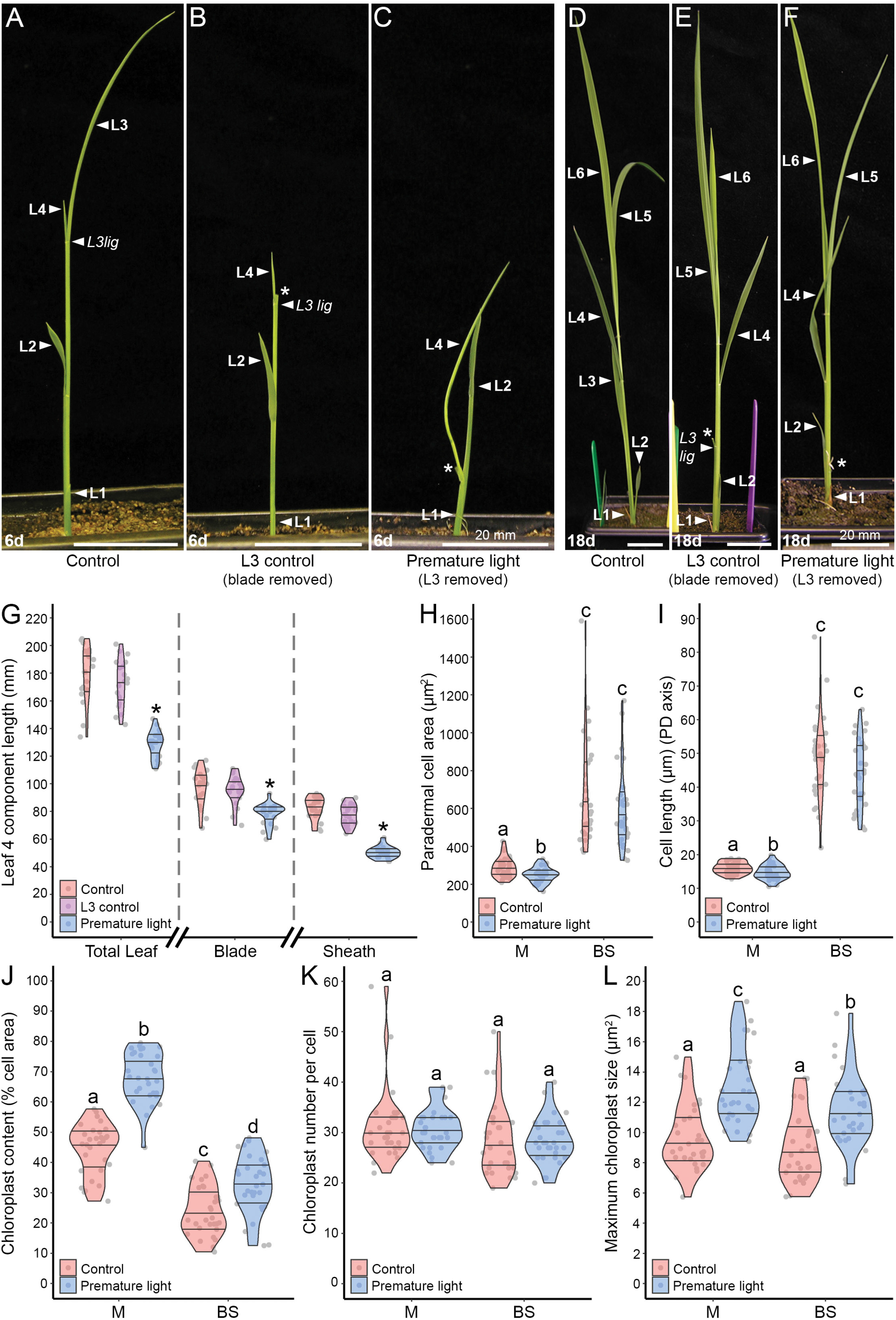
Premature exposure of leaf 4 primordia to light reduces both final leaf 4 length and expansion of mesophyll but not bundle sheath cells. **A-F**. Premature light exposure treatments, when applied to leaf 4 at emergence (**A**, control; **B**, leaf 3 blade only removed; **C**, leaf 3 removed exposing leaf 4 primordium) and subsequent whole plant phenotypes at leaf 4 maturity (**D**, control; **E**, leaf 3 blade only removed; **F**, leaf 3 removed). The positions of leaves 1-6 (L1 – L6) and the ligule of leaf 3 (L3 lig) are indicated by arrowheads. Asterisks denote the point at which leaf 3 has been cut. **G**. Comparison of leaf 4 length at maturity between control, leaf 3 control and early light treatments showing the total length, blade and sheath lengths of leaf 4. Violin plots represent data distribution density, with their width corresponding to frequency of datapoints at that value. Median value and upper and lower quartiles are represented by the middle, upper and lower horizontal lines, respectively. Asterisks denote a significant difference (p < 0.05) between treatments within a particular measurement category. *n* = 20. **H-L**. Comparison of the paradermal cell area (**H**), cell length (**I**), relative chloroplast content (**J**), chloroplast number (**K**) and chloroplast size (**L**) of mesophyll (M) and bundle sheath (BS) cells between control and early light treatments. Violin plots represent data distribution density, with their width corresponding to frequency of datapoints at that value. Median value and upper and lower quartiles are represented by the middle, upper and lower horizontal lines, respectively. Cell type + treatment combinations denoted with different letters within each plot are significantly different from each other (p < 0.05). Comparisons within all characters were made using transformed datasets to meet the assumptions of statistical tests (see **Supplementary Dataset S4**). *n* = 30 (10 cells of each type measured from 3 individual plants per treatment).

To provide insight into how premature exposure to light reduced leaf length, the leaf 4 primordium was prematurely exposed to light (**Fig 4C**) and cell morphology at the base of the blade examined after 3 days. This stage was selected to avoid the masking of any effects of light on mesophyll cell development due to chloroplast crowding. In control plants this blade tissue was not fully exposed to light because it was still surrounded by the leaf 3 sheath but cell expansion was already complete (**Supplementary Fig. S5A, B**). Whilst premature light reduced the area and length of mesophyll cells (p < 0.05; **Fig. 4H, I**) no statistically significant effects were detected on bundle sheath cell area or length (p > 0.05). Much like the mesophyll, epidermal cells exhibited a significant reduction (p < 0.05) in area and length when prematurely exposed to light (**Supplementary Fig. S5C-E**). The proportion of cell occupied by chloroplast was significantly increased in prematurely exposed mesophyll and bundle sheath tissue compared with cells not exposed to light (p < 0.05, **Fig. 4J**) demonstrating that both cell types had perceived the light signal. This was not associated with an increase in chloroplast number (p > 0.05, **Fig. 4K**) but instead an increase in chloroplast size (p < 0.05, **Fig. 4L**). These results suggest that the cell types responded differently to a premature light signal, with cell expansion being inhibited in the epidermis and mesophyll layers but not in the bundle sheath. Consistent with the hypothesis that light signalling promotes exit from the cell cycle, early light reduced leaf length. As the bundle sheath is continuous along the leaf this indicates that in this tissue light must have stopped cell divisions earlier, or that divisions occurred less frequently.

In summary, we provide evidence that bundle sheath and mesophyll cells produce chloroplast compartments of similar size, but continued expansion of the bundle sheath cells leads to a lower percentage chloroplast occupancy. The relationship between bundle sheath cell size and chloroplast content was significantly influenced by cytokinin and gibberellin. Preventing or delaying exposure to light increased total leaf length but decreased length of its constituent cells indicating that light signalling regulates cell division during leaf development. Interestingly, bundle sheath cells were still able to expand in response to a late light signal, resulting in a change in cell shape at maturity. Finally, the notion that light regulates the cell cycle in rice leaves is supported by an orthogonal experiment in which premature exposure to light reduced final leaf length but did not change bundle sheath cell length. Our data are therefore consistent with light playing an antagonistic role in the control of bundle sheath cell division.

## DISCUSSION

### Chloroplast biogenesis in rice bundle sheath cells responds to cell size and this relationship is influenced by hormone signalling

Although it is well known that cell and chloroplast development influence each other during leaf development such that individual cell types achieve set levels of chloroplast, and different cell types contain differently sized chloroplast compartments, the mechanisms that co-ordinate these processes are unclear. The rice bundle sheath has previously been shown to contain less chloroplast compared with mesophyll cells (Wang *et al*., 2017b). This is consistent with observations in many other C_3_ species (Williams *et al*., 1989, Kinsman and Pyke, 1998; McKown and Dengler, 2007; Khoshravesh *et al*., 2016, 2020). Here, we show that both chloroplast size and number are reduced in mature bundle sheath cells compared with the mesophyll, suggesting that chloroplast biogenesis is partially repressed. Importantly, regression analysis demonstrated that, as with mesophyll cells, chloroplast content in the bundle sheath is cell-size dependent: as paradermal cell area increased so did total chloroplast area per cell. However, the rate of chloroplast biogenesis was not sufficient to keep pace with cell expansion, and this resulted in a negative relationship between relative chloroplast content and cell size. Chloroplast number but not size changed with increasing bundle sheath cell size, a response previously observed in the mesophyll cells of both *A. thaliana* and wheat (Ellis and Leech, 1985; Pyke and Leech, 1991). In *A. thaliana* altered chloroplast number has been shown to be an outcome rather than a driver of increased chloroplast content. For example, in mutants with impaired plastid division total chloroplast area still correlated tightly with cell size (Pyke and Leech, 1992) but individual chloroplast size was increased (Pyke *et al*., 1994; Miyagishima *et al*., 2006; Schmitz *et al*., 2009) Conversely, over-expression of the *Plastid Division Genes PDV1* and *PDV2* resulted in more numerous but smaller chloroplasts (Okazaki *et al*., 2009).

These publications and the analysis reported here used two-dimensional imaging of three-dimensional cells. Thus, information relating to cell depth and volume is not captured. For processes relating to the capture of light and rates of photosynthesis expressed per unit leaf area, such two-dimensional analysis would seem appropriate as light is intercepted on a leaf area basis. However, processes directly affected by volume such as diffusion of gases and metabolites are more difficult to interpret without three dimensional analyses. Most notably, two-dimensional imaging is likely to under-report the volume and number of chloroplasts in mesophyll cells where they are more closely packed, and uniformly over-estimate the relative chloroplast content of cells compared with their relative volume due to cell depth being discounted. To maximise the accuracy of chloroplast measurements and counts, in this study individual cells were each imaged at multiple focal planes (see Materials and Methods). The conclusion from our two-dimensional analysis that chloroplast number but not size responds to cell size matches that of previous studies as described above, and using geometric extrapolation to estimate bundle sheath cell volumes further supports a positive relationship with chloroplast number. We detected the same cell-size responses in chloroplast biogenesis in both rice bundle sheath and mesophyll cells, and as such conjecture that a similar mechanism regulates this response in both cell types. If so, the reduced relative chloroplast content of the bundle sheath (and the enhanced chloroplast content of C_4_ bundle sheath cells) could arise through differential regulation of such a conserved cell-size dependent network.

The hormones auxin, CK and GA have each been linked to aspects of cell and chloroplast development. For example, in *A. thaliana* auxin regulates cell division and final organ size in shoots (Hu *et al*., 2003; Schruff *et al*., 2006) and has been shown to limit root cell expansion (Löfke *et al*., 2015). CK promotes multiple aspects of chloroplast development including chloroplast division (Cortleven and Schmülling, 2015) and GA regulates expansion and division of leaf epidermal cells (Achard *et al*., 2009; Claeys *et al*., 2012) with some evidence that it also regulates chloroplast division in *A. thaliana* and rice (Jiang *et al*., 2012). In fact, the density of chloroplasts per cell area was increased in extremely GA-deficient mutants, but a strong linear relationship between cell size and chloroplast number was retained (Jiang *et al*., 2012). The extent to which GA signalling directly regulates this relationship or whether this was an indirect consequence of impaired cell expansion is not clear. We found that exogenous treatment with these hormones led to the relationship between relative chloroplast content and bundle sheath cell size became significantly more positive, and the relationship between total chloroplast area and cell size was also enhanced with CK and GA treatment. As mean cell size was unaffected, this indicates enhanced chloroplast biogenesis. We note that although no effect of hormone treatment was detectable in mesophyll cells, their high chloroplast content could mask or prevent such an increase being easily detected. In *A. thaliana* these hormones are part of a chloroplast regulatory network that includes the *GLK* gene family (Wang *et al*., 2017a). Moreover, in rice overexpression of *GLK* can drive increased chloroplast content in the bundle sheath (Wang *et al*., 2017b). Our analysis therefore suggests that the mechanism for cell-size-dependent regulation of chloroplast biogenesis either involves or is itself regulated by CK and GA.

### Light signalling promotes exit of the rice bundle sheath from cell division

In contrast to dicotyledonous models such as *A. thaliana*, leaves of monocotyledonous plants develop via a basal zone of cell proliferation from which cells emerge, cease dividing and then expand as they progress distally (Conklin *et al*., 2019). Here we report that when rice was grown in the dark leaf length at maturity was increased whereas bundle sheath and mesophyll lengths in the same leaves decreased, indicating that increased leaf length was achieved through prolonged cell division. This interpretation is consistent with the changing distributions of bundle sheath cell lengths, where dark-grown plants lacked a population of the longest cells. Consistent with light signalling promoting exit of *A. thaliana* leaf epidermal cells from cell division via chloroplast retrograde signalling (Andriankaja *et al*., 2012), developing rice leaves exposed prematurely to light were shorter at maturity, but bundle sheath cell length was unchanged. Conversely, GA treatment apparently prolonged cell division. This is consistent with a kinematic analysis of maize leaf development which found that exit from the cell-cycle and maturation was accelerated in the loss-of-function GA biosynthesis mutant *dwarf3* and delayed in GA-overexpressing transgenic lines, whilst final cell length was unaffected in both (Sprangers *et al*., 2020). The lack of an effect of GA treatment on final cell size, whilst surprising, might therefore reflect a difference between leaf development in monocotyledons and dicotyledons. GA and light signals interact in circumstances such as photomorphogenesis of dark-grown seedlings (Mazzella *et al*., 2014) with GA levels reduced by exposure to light (Reid *et al*., 2002). Increased cell division in dark-grown leaves might therefore be explained through increased GA, but the reduced length of dark-grown bundle sheath cells also suggests additional pathways. Surprisingly, the length of both mock and GA-treated cells grown under light from leaf 4 resembled dark-grown cells in leaf 2 rather than light-grown cells (**Supplementary Dataset S3**). This could be due to the different leaf position analysed between these experiments, as variation in mature bundle sheath volume have previously been recorded between leaves 4 and 7 (Wang *et al*., 2017b). Our findings thus strongly support a role for light signalling in promoting exit of the bundle sheath from the cell cycle, which may act in part through an established interaction with GA biosynthesis.

Although evolutionary trajectories attempting to rationalise emergence of C_4_ anatomy have been proposed from analysis of C_3_, C_3_-C_4_ intermediate, and C_4_ leaves (McKown and Dengler, 2007; Muhaidat *et al*., 2011; Christin *et al*., 2013; Williams *et al*., 2013; Lundgren *et al*., 2014; Khoshravesh *et al*., 2016; Reeves *et al*., 2018; Lundgren *et al*., 2019; Khoshravesh *et al*., 2020) the specific genetic changes through which these adaptations arose remain less well understood. Analysis of the evolution of the C_4_ biochemical pathway so far indicate that changes arose by modifying pre-existing ancestral *cis*-elements and *trans-*factors (Matsuoka *et al*., 1993,1994; Brown *et al*., 2011; Kajala *et al*., 2012; Williams *et al*., 2016; Reyna-Llorens *et al*., 2018). The evolution of C_4_ photosynthesis is frequently accompanied by a change in the shape of bundle sheath cells, such that they become shorter and wider (McKown and Dengler, 2007; Muhaidat *et al*., 2007; Khoshravesh *et al*., 2020). We found that in rice leaves grown entirely in the absence of light bundle sheath cells were shorter but also less wide than light-grown controls, whereas bundle sheath cells exposed to delayed light matured to a similar width as light-grown controls but obtained a final length similar to dark-grown cells. From this, we infer that light signalling has a separable role promoting expansion of the bundle sheath and that whilst expansion along the proximodistal axis is constrained through physical attachment to vascular tissues lateral expansion is not constrained to the same extent. These observations indicate that the unique morphology of bundle sheath cells from C_4_ species could have evolved through changes in their developmental responses to light.

A transgenic cell division reporter in *A. thaliana* further supports underlying differences in bundle sheath and mesophyll development, with cell division evident in vasculature after it has finished in the mesophyll or epidermis (Donnelly *et al*., 1999). In C_4_ maize, light-dependent gene regulatory networks have been shown to differ between bundle sheath and mesophyll cells, such that they exhibit differential transcriptional responses to blue and red light (Hendron and Kelly, 2020) and it has been suggested that these differences may have contributed to the evolution of the C_4_ bundle sheath. We therefore propose that bundle sheath and mesophyll cells of C_3_ plants possess differences in the underlying genetic networks underpinning responses to light, and that adjustments to the existing network in bundle sheath cells modifies their morphology to allow the evolution of C_4_ photosynthesis.

## Abbreviations

BAP: 6-benzylaminopurine
BS: bundle sheath
C_3_: 3-carbon
C_4_: 4-carbon
CK: cytokinin
GA: gibberellic acid
M: mesophyll
ML: mediolateral
NAA: 1-Napthaleneacetic acid
PD: proximodistal
RuBisCO: Ribulose Bisphosphate Carboxylase Oxygenase

## Supplementary Data

The following supplementary data are available at JXB online.

**Figure S1**. Plant-to-plant variation in bundle sheath and mesophyll cell characters.

**Figure S2**. Geometric estimation of bundle sheath cell volumes.

**Figure S3**. Hormone treatments significantly affect relative and absolute chloroplast content in rice bundle sheath but not mesophyll.

**Figure S4**. Rice leaf growth responses to altered light exposure.

**Figure S5**. Comparison of leaf 4 development in premature light treatment experiments against normal leaf 4 development.

**Dataset S1**. Statistical analysis and raw data of mature leaf 4 BS and M cell characteristics.

**Dataset S2**. Statistical analysis and raw data of leaf 4 BS and M cell phenotypes in response to auxin, BAP or GA_3_ treatment.

**Dataset S3**. Statistical analysis and raw data of the effect of delayed light exposure on the development of leaf 2 BS and M cells.

**Dataset S4**. Statistical analysis and raw data of the effect of early light exposure on the development of leaf 4 BS, M and E cells.

## Acknowledgements

We are grateful to Dr. Roxana Khoshravesh for the single cell isolation protocol and for discussions on optimisation in rice. We are grateful to staff at the University of Cambridge Plant Growth Facility and Susan Stanley for support in growing plant material.

## Author contributions

ARGP and JMH: conceptualisation; ARGP: formal analysis; JMH: funding acquisition; ARGP: investigation; JMH: supervision; ARGP: writing-original draft preparation: ARGP and JMH: writing-review and editing.

## Conflicts of interest

The authors declare that they have no conflicts of interest associated with this work.

## Funding

This work was supported by BBSRC Grant BBP0031171 to JMH, and the C_4_ Rice project grant from The Bill and Melinda Gates Foundation to the University of Oxford (2015–2019). For the purpose of open access, the authors have applied a Creative Commons Attribution (CC BY) license to any Author Accepted Manuscript version arising from this submission.

## Data availability

All data supporting the findings of this study are available within the paper and within its supplementary materials published online.

## Notes

### Competing Interest Statement

The authors have declared no competing interest.

