## Supplementary material for "Delayed exposure of rice to light partially phenocopies C_4_ bundle sheath characteristics by reducing cell length": S Data Figs

**SUPPLEMENTARY FIGURES S1-S5**

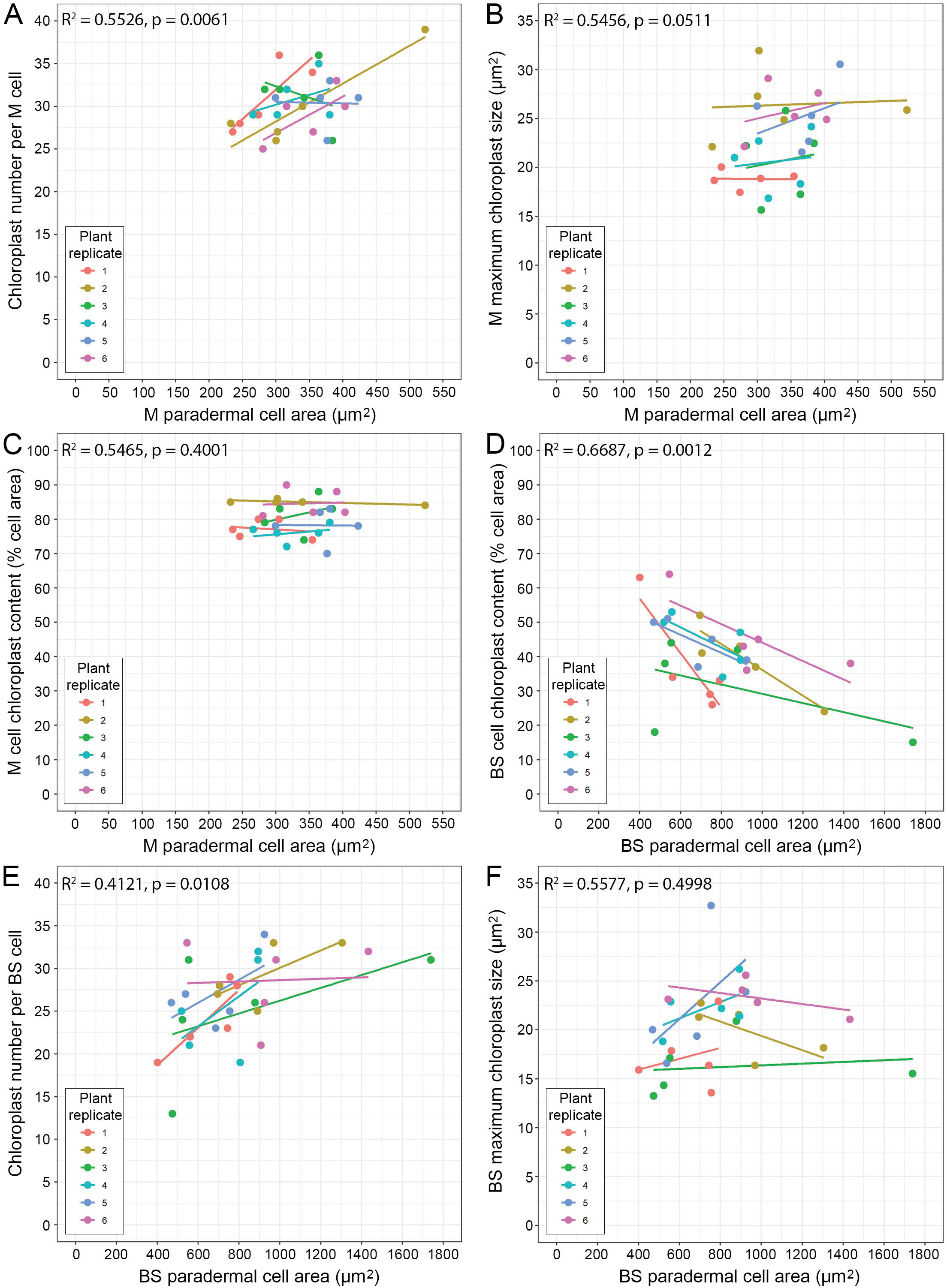

**Supplementary Fig. S1.** Plant-to-plant variation in bundle sheath and mesophyll cell characters.

**A, B.** Linear regression analyses of chloroplast number (**A**) and size (**B**) in mature mesophyll cells isolated from leaf 4, including the plant from which cells were isolated as a model parameter.

**C, D.** Linear regression analyses of relative chloroplast size content against cell area in mature mesophyll (**C**) and bundle sheath cells (**D**) isolated from leaf 4, including the plant from which cells were isolated as a model parameter.

**E, F.** Linear regression analyses of chloroplast number (**E**) and size (**F**) in mature bundle sheath cells isolated from leaf 4, including the plant from which cells were isolated as a model parameter.

BS, bundle sheath; M, mesophyll. *n* = 30 (5 cells measured from 6 individual plants). P-values shown denote the significance of cell area as a factor in determining the response variate shown (Y axis). Full linear regression models and associated p-values are provided in **Supplementary Dataset S1**.

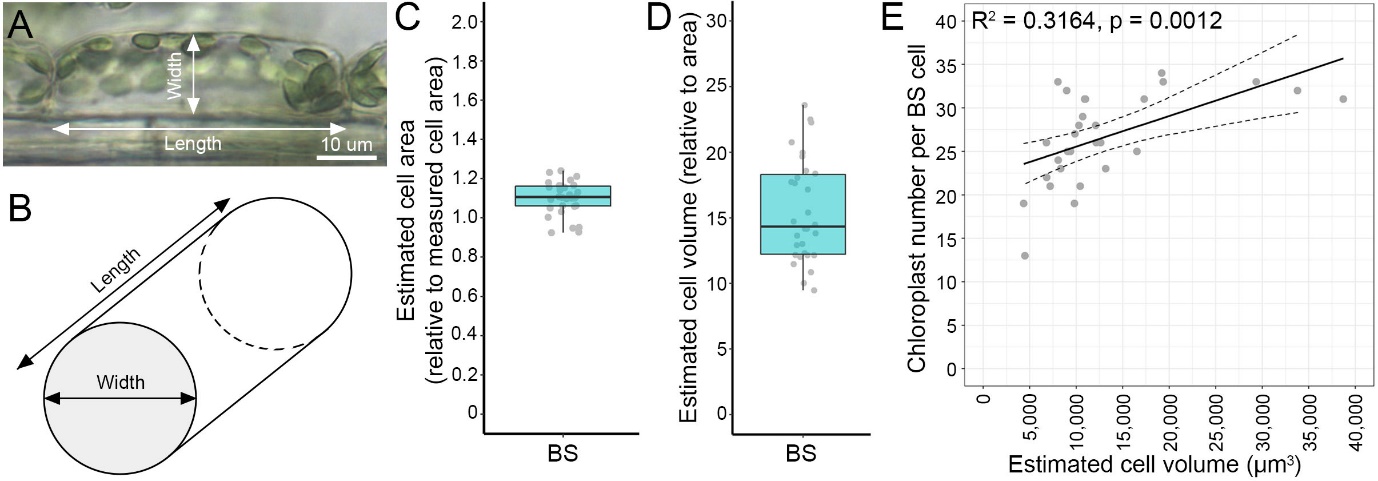

**Supplementary Fig. S2.** Geometric estimation of bundle sheath cell volumes

**A-B.** Linear measurements of cell length and cell width from mature bundle sheath cells (**A**) were used to calculate individual cylinder volumes (**B**) using the formula V = (Pi*(^d^/_2_))^^2^ * L, taking cell width as the circle diameter (d) and cell length as the cylinder length (L). The irregular shape of mesophyll cells (**Fig. 1A**) meant that 3D volumes could not be similarly estimated for this cell type.

**C.** The accuracy of geometric estimation approach was tested by comparing the paradermal cell area estimated from linear measurements (calculated by multiplying cell length by cell width) against the corresponding cell area measured directly for each cell. Calculated cell area values are shown relative to directly-measured cell area. The mean ratio of calculated-to-measured cell areas was 1.09 and significantly different from 1 (two-tailed one-sample T-test, p < 0.0001), indicating that estimating cell area from linear measurements over-estimates individual cell area by approximately 9%.

**D.** The mathematical independence of estimated 3D cell volumes from 2D cell area measurements was confirmed by plotting estimated cell volumes relative to their corresponding measured cell areas. Datapoints were found to vary between an increase of approximately 10 to 25-fold, demonstrating that 3D volume estimations do not represent a simple mathematical transformation of 2D data. This is because cell volumes are calculated from two independent values (cell length and cell width).

**E.** Linear regression analysis of chloroplast number per bundle sheath cell against corresponding estimated cell volume. P-value shown denotes the significance of cell volume as an explanatory factor. R^2^ values indicate the explanatory power of the fitted model.

BS, bundle sheath. *n* = 30 (5 cells measured from 6 individual plants). Full linear regression models, associated p-values and geometric estimates of cell area and volume are provided in **Supplementary Dataset S1**.

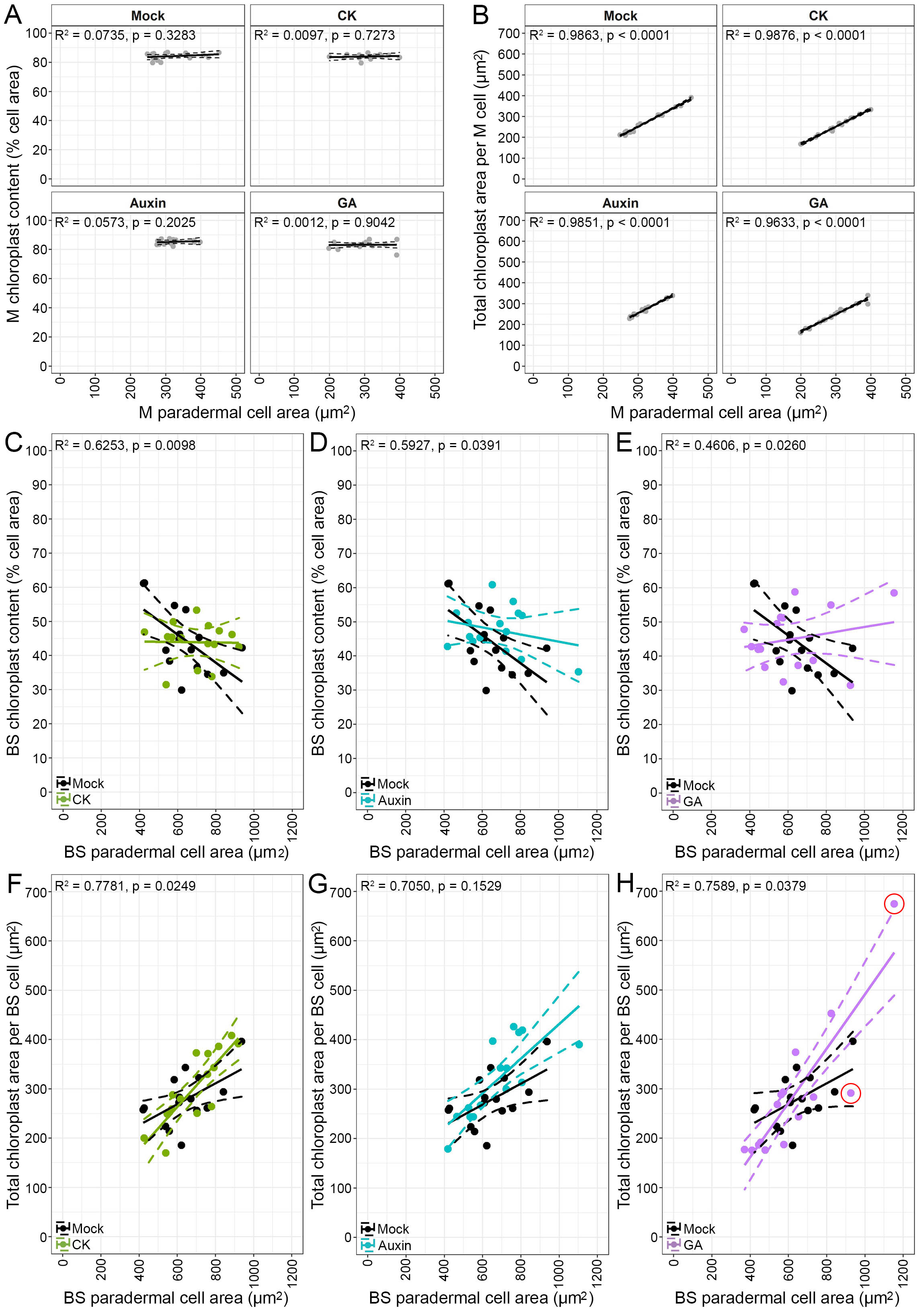

**Supplementary Fig. S3.** Hormone treatments significantly affect relative and absolute chloroplast content in rice bundle sheath but not mesophyll.

**A**-**B**. Regression analyses of relative chloroplast content (**A**) and total chloroplast area per cell (**B**) against cell area in mesophyll cells under mock (0.1% EtOH v/v), cytokinin (5 µM BAP), auxin (5 µM NAA) and gibberellin (100 µM GA_3_) treatments, showing the line of best fit (solid line) and 95% confidence interval (dashed line) of the model fitted to the data (grey circles). R^2^ and p-values and are from regression analyses of the data against cell area within each treatment, denoting the explanatory power of the fitted model and the significance of cell area as an explanatory variate within it.

**C**-**H**. Linear regression analyses of relative chloroplast content (**C-E**) and absolute chloroplast area per cell (**F-H**) against cell area in mature bundle sheath cells, comparing mock treatment (0.1% EtOH v/v) against cytokinin (5 µM BAP) (**C,F**), auxin (5 µM NAA) (**D,G**) and gibberellin (100 µM GA_3_) (**E,H**), respectively. P-values shown denote the significance of the interaction between hormone treatment and cell area. R^2^ values indicate the explanatory power of the fitted model. Red circles in **H** denote outlier datapoints identified by analysis of residuals as potentially having an unduly high impact on the regression analysis. This analysis was also performed with those datapoints excluded, with no change in the significance of the p-value obtained (**Supplementary Dataset S2**).

BS, bundle sheath; M, mesophyll. *n* = 15 (5 cells measured from 3 individual plants). Full linear regression models and associated p-values are provided in **Supplementary Dataset S2**.

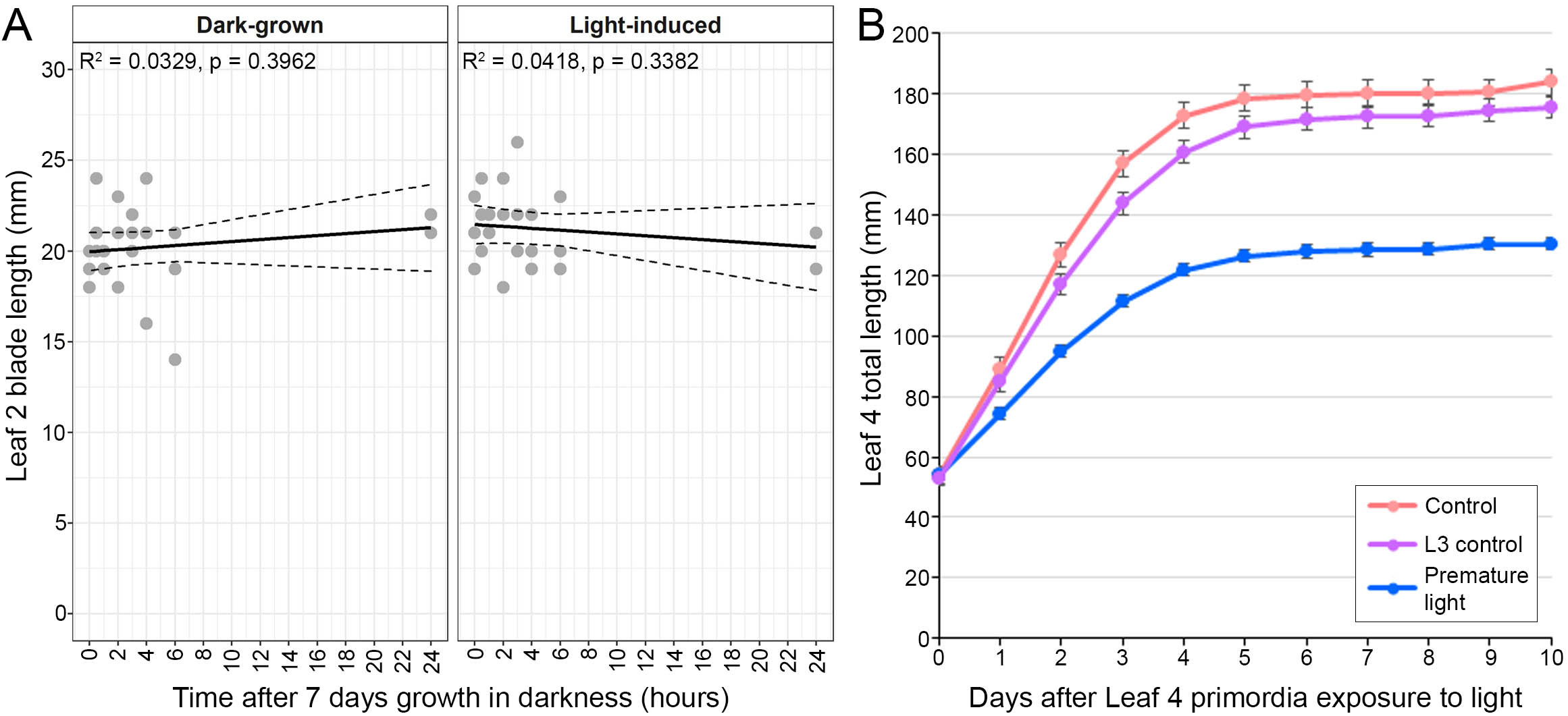

**Supplementary Fig. S4.** Rice leaf growth responses to altered light exposure

**A.** Regression analyses of dark-grown leaf 2 blade length against time after exposure to light at seven days after germination or continued growth in the dark (as shown), showing the line of best fit and 95% confidence interval of the model fitted to the data (grey circles). P-values and R^2^ values are from regression analyses of the data against time within each light treatment, denoting the significance of time as an explanatory variate of leaf 2 length and the explanatory power of the fitted model, respectively. Independent plants were sampled at each timepoint due to the destructive nature of the sampling. Leaf 2 length was not found to increase with time after 7d growth in the dark. *n* = 24. All raw data and full regression analyses are given in **Supplementary Dataset S3**.

**B.** Growth response of leaf 4 to premature light (entire leaf 3 removal at L4 primordium emergence) and leaf 3 blade removal (**Fig. 3A-C**) over time. Values shown are the mean total leaf length ± SE, with repeated daily measurements taken of the same plants. *n* = 20. All raw data are given in **Supplementary Dataset S4**.

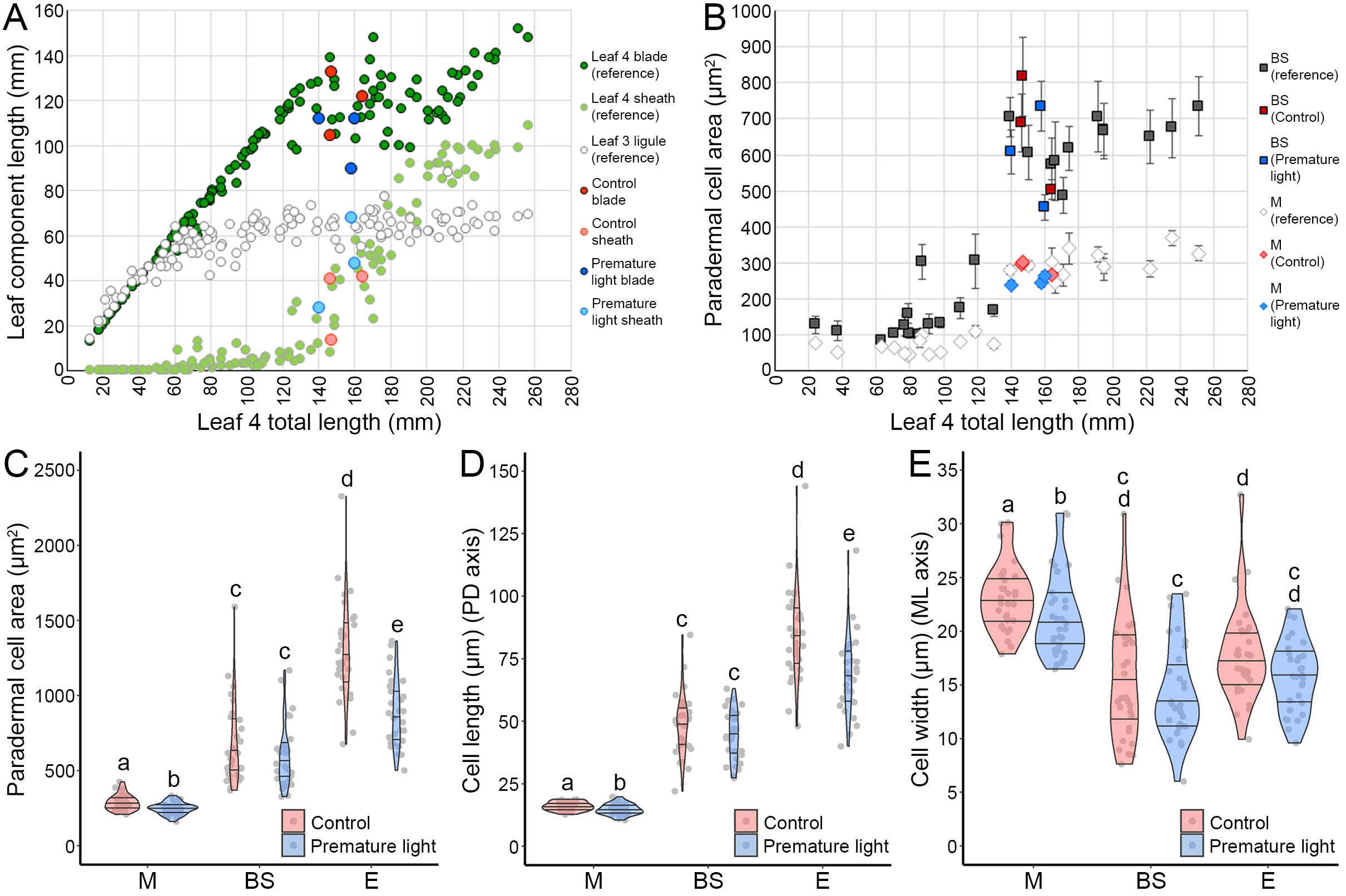

**Supplementary Fig. S5.** Comparison of leaf 4 development in premature light treatment experiments against normal leaf 4 development.

**A**, Lengths of leaf 4 components (blade and sheath) and the leaf 3 sheath enclosing leaf 4 sampled from 130 reference plants plotted against total leaf 4 length at harvesting (individual datapoints shown). Leaf length measurements from control and premature light-treated plants sampled in cell response experiments (**Fig. 4H-L**) are shown in relation to this gradient (blue and yellow datapoints, respectively). All reference leaf length measurements are provided in **Supplementary Dataset S4**.

**B.** Paradermal cell area measurements of mesophyll and bundle sheath cells from the basal 5 mm of the leaf 4 blade across the reference gradient of total leaf 4 lengths. Values shown are the mean cell area ± SE from 5 cells isolated from each reference plant. Cell area measurements from control and premature light-treated plants sampled from the same tissue (**Fig. 4H**) are included for comparison. All reference cell measurements are provided in **Supplementary Dataset S4**.

**C-E.** Comparison of the effect of control and premature light treatments on leaf 4 paradermal epidermal cell area (**C**), cell length (**D**, leaf proximodistal axis) and cell width (**E**, leaf mediolateral axis). Mesophyll and bundle sheath cell data (**Fig. 4H-I**) is reproduced here for ease of comparison. Violin plots represent data distribution density, with their width corresponding to frequency of datapoints at that value. Median value and upper and lower quartiles are represented by the middle, upper and lower horizontal lines, respectively. Cell type + treatment combinations denoted with different letters within each plot are significantly different from each other (p < 0.05). BS, bundle sheath; E, epidermis; M, mesophyll. *n* = 30 (10 cells of each type measured from 3 individual plants per treatment). All cell measurements are provided in **Supplementary Dataset S4**.
